# Low Physiological Arousal in Mental Fatigue: Analysis of Heart Rate Variability during Time-on-task, Recovery, and Reactivity

**DOI:** 10.1101/2020.08.24.264812

**Authors:** András Matuz, Dimitri van der Linden, Zsolt Kisander, István Hernádi, Karádi Kázmér, Árpád Csathó

## Abstract

Heart Rate Variability (HRV) has been suggested as a useful tool to assess fatigue-sensitive psychological operations. The present study uses between and within-subject design to examine the temporal profile of HRV including the changes related to reactivity, time-on-task (ToT), and recovery on a cognitively demanding task. In the fatigue group, participants worked on a bimodal 2-back task with a game-like character (the gatekeeper task) for about 1.5 hours, followed by a 12-minute break, and a post-break block of performance (about 18 min). In the control group, participants watched documentaries. We hypothesized that mental fatigue is associated with low physiological arousal and increasing vagal-mediated HRV as a function of ToT. We also analysed the trial-based post-response cardiac activity as a physiological indicator of task-related motivation. Relative to the control, ToT was associated with an elevated level of subjective fatigue, decreased heart rate, and increased HRV most robustly in the vagal-mediated components. Based on fatigued participants’ post-error cardiac slowing, and post-error reaction time analyses, we found no evidence for motivation deficit in association with increasing HRV and ToT. The present findings support the low arousal state of mental fatigue and suggest that primarily the vagal components of the HRV spectrum are indicative of fatigue. In addition, the study provides evidence that many HRV indices might be changed not only in a fatiguing condition but also if individuals are engaged in a prolonged non-fatiguing activity. This finding emphasizes the relevance of control conditions in ToT experiments.

## Introduction

Mental fatigue has a detrimental effect on a wide range of prefrontal-cortex loaded cognitive functions, reduces the willingness to exert further effort, and is frequently accompanied with reduced performance efficiency and an increased number of errors (e.g. Ackerman, 2011; Lorist et al., 2005; Van der Linden, 2011). Fatigue and its reduced task-specific motivation are often suggested to be accompanied with a decreased physiological arousal level (Van der Linden, 2011; Hopstaken et al., 2015a). From this perspective, it is understandable why previous studies have explored the association between mental fatigue and heart rate variability (HRV) as an index of cardiac autonomic regulation (i.e. the variability in intervals between successive heartbeats). Those studies converged on the conclusion that HRV is a significant marker of fatigue that can predict the concurrent drop in cognitive performance due to time-on-task (ToT; see e.g. Gergelyfi et al., 2015; Mizuno et al., 2014; Mizuno et al., 2011; Tran et al., 2009; Segerstrom and Nes, 2007; Fairclough and Houston, 2004).

Nevertheless, as fatigue is a complex, multifaceted state and HRV has many calculable components that have diverse sources, this greatly complicates the exact biopsychological interpretation of the HRV – fatigue associations. Therefore, in recent studies, it has been emphasized that for better comparability, future studies should build on more recent insights regarding the physiological and methodological substrates of HRV (Sassi et al., 2015; Laborde et al., 2017). Regarding this, vagal control on cardiac activity, and the associated HRV level is known to be strongly determined by the activity of the locus coeruleus-norepinephrine system (LC-NE system; Mather et al., 2017), which has been suggested to be less active under fatigue (Hopstaken, et al., 2015a, 2015b; Van der Linden, 2011). Given the above recommendations and the inhibitory activity of LC-NE on vagal activity, in the present study, we hypothesized that primarily the vagal components of HRV are predictive for fatigue-related changes, that is, the root mean square of successive differences (RMSSD), pNN50, and the high frequency (HF) component of HRV. RMSSD and pNN50 are the primary time-domain measures used to estimate vagal-mediated changes and are relatively free of respiration signal components (Thayer and Lane, 2000; Malik, 1996). The HF component also reflects vagal tone, but it more strongly corresponds to the respiratory cycle (Malik, 1996; Laborde et al., 2017).

In a recent study, we found evidence for increased vagal-mediated HRV with increasing ToT on a bimodal task-switching task (Matuz et al., 2019). This finding provided further support for the notion that prolonged task performance may be associated with reduced activity of the LC-NE system and a lowered arousal state. An important limitation of that HRV measurement, however, was the lack of comparison with a non-fatiguing condition. Therefore, in the current study, we compared the HRV functions in two different groups: a group in which participants have to engage in a cognitively demanding task (i.e. a bimodal 2-back task; Heathcote et al., 2015; Heathcote et al., 2014) and a non-fatigue group in which participants watch documentary films. In the fatigue group, we hypothesized an increasing HRV (i.e. a lowered physiological arousal) as a function of ToT. In contrast, we expected to find no change in HRV in the control, documentary viewing group.

In designing the study, we followed the recent recommendation of Laborde and colleagues (2017) by testing the changes in HRV in the resting and active phases of the experiment using the “three Rs” concept: resting, reactivity, and recovery. Specifically, in addition to the change in HRV while performing the task for a prolonged period (i.e. time-on-task), we also explored the change from active task performance to a resting state (recovery) and, conversely, from a resting state to active performance (reactivity). In order to assess the recovery-related effects, the last block of trials in our study was preceded by a break period of 12 minutes. Reactivity and recovery-related changes in HRV can be interpreted within a stressor-strain model (Sonnentag, 2011). As Sonnentag (2011) stated, the physiological changes in reactivity refers to the immediate physiological needs when the stressor is present. Conversely, recovery refers to all of those physiological processes that occur when the stressor disappears and the system tends to return to its baseline, prestressor level. This suggests that by investigating the reactivity and recovery related changes in HRV we can identify more clearly the arousal regulatory mechanisms in an initial task performance phase and in a phase when individuals became fatigued after prolonged performance.

We have referred to the possibility that the nature of fatigue-related arousal changes may depend on the person’s motivational stance – that is, whether or not they are still motivated to perform well. Therefore, to gain insight into the participants’ motivational stance, we also explored the changes in phasic cardiac activity (i.e. heart rate change) after the participants’ responses. Specifically, several lines of evidence suggest that phasic heart rate deceleration after an inaccurate response might be a physiological correlate of general performance monitoring and error awareness (Hajcak et al., 2004). Performance monitoring has been found to show a decline with increasing mental fatigue in relation to individuals’ decreased motivation to perform the task (Boksem et al., 2006). These earlier findings underline the relevance of analysing changes in post-response phasic cardiac activity in the current study. Importantly, to our knowledge, no previous study has addressed post-response cardiac activity in fatigue research.

To summarize, in the present study, we aimed to examine the temporal profile of HRV, including the changes related to ToT, recovery, and reactivity. We hypothesized that the state of mental fatigue is accompanied with a low physiological arousal state, reflecting increased vagal-mediated HRV as a function of time-on-task. We also argue, however, that the examination of reactivity-and recovery-related changes are important to draw more robust conclusions about how physiological arousal mechanisms control the performance of a fatiguing, cognitively demanding task. In addition we analysed the trial-based post-response cardiac activity as a physiological indicator of task-related motivation.

## Materials and Methods

### Participants

Forty-four participants (under-and post-graduate students), in a medication-free health condition, with normal hearing and normal or corrected-to-normal vision participated in the study; there were 22 participants in the fatigue group (i.e. gatekeeper task) and 22 participants in the non-fatigue control (i.e. documentary-viewing) group. Because of technical failures, the data of three participants were excluded from the analysis. Thus, the final dataset used for the analyses contained data from 20 participants (11 females, mean age: 21.2 with SD of 2.21, range: 19-27) in the fatigue group and 21 participants (11 females, mean age: 22.5 with SD of 3.9, range: 18-29) in the control group. Participants in the two groups were matched in age (t(39) = −1.33, *p* = .19) and gender (χ^2^ = .22, *p* = .64). All participants provided written consent.

The minimum sample size to ensure the statistical power of the main effect of ToT on HRV were estimated based on our recent study (Matuz et al., 2019), as well as other studies that were published recently and examined either the difference between an active task-performance and a resting period or the modulation of HRV by a prolonged task performance (e.g. Hidalgo-Muñoz et al., 2018; Delliaux etal., 2019; Gergelyfi et al., 2015; Pendleton et al., 2016). By applying the lowest effect size for two groups (by Gpower 3.1., Faul et al., 2007), the recommended minimum sample size per group was 18 participants to achieve a power level of 90% and alpha < 0.05. Thus, the final sample of 20 and 21 participants, respectively, had the appropriate statistical power to detect the effects we aimed to examine.

### Task and Stimuli

#### Fatigue group - Gatekeeper task

Participants in the fatigue group performed an adapted version of the gatekeeper task (Heathcote et al., 2014, 2015) which is a dual 2-back task with visual and auditory stimuli. The gatekeeper task has a game-like character: participants need to imagine that they are a nightclub doorperson and need to memorize the door and the password used by the guests of the club for entry. This game-like feature of the task is an asset because it is expected to enhance task engagement, which may lead to less boredom during ToT. In each trial, the visual stimuli (i.e. three door images, one of which is highlighted in red) and the auditory stimulus (i.e. one spoken letter) were presented simultaneously (see Figure 1). Four different stimulus conditions were prepared. For dual target condition, both the visual and auditory stimuli were identical to those presented two trials earlier. For the single target conditions, a 2-back match occurred either for the auditory stimulus (single auditory target condition) or for the visual stimulus (single visual target condition). For the no target condition, both the visual and the auditory stimuli were different to the stimuli shown two trials earlier. In each trial, participants were instructed to indicate by pressing a key whether there is a 2-back match in any modality, which in this case would imply that the ‘guest’ would not be allowed to enter the night club. In the instructions, it was emphasized that both speed and accuracy are equally important. A new trial began after a 2.5s interval after response.

**Figure 1.**
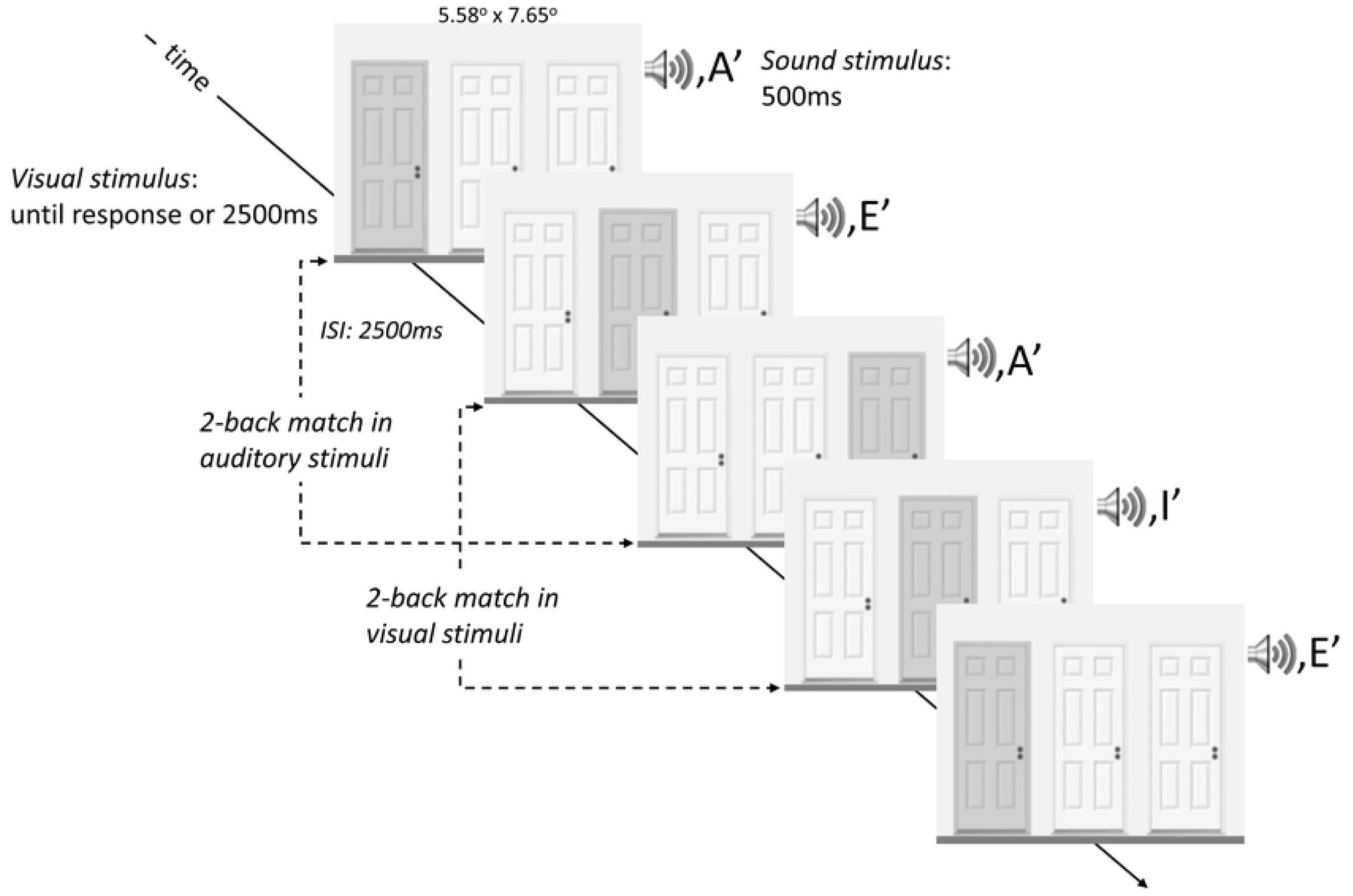
Schematized sequence of trials in the gatekeeper task. The gatekeeper task is a dual 2-back task with visual and auditory stimuli. The gatekeeper task has a game-like character: participants need to imagine that they are a nightclub doorperson and need to memorize the door and the password used by the guests of the club for entry. In each trial, the visual stimuli (i.e. three door images, one of which is highlighted in red) and the auditory stimulus (i.e. one spoken letter) were presented simultaneously. For dual target condition, both the visual and auditory stimuli were identical to those presented two trials earlier. For the single target conditions, a 2-back match occurred either for the auditory stimulus (single auditory target condition) or for the visual stimulus (single visual target condition). For the no target condition, both the visual and the auditory stimuli were different to the stimuli shown two trials earlier. In each trial, participants were instructed to indicate by pressing a key whether there is a 2-back match in any modality.

#### Control group: Documentary film watching

The participants in the control group watched three documentary films (about 30 minutes each) for 1.5 hours: Planet Earth Episode 7 Great plains (2007); When we left Earth – The NASA missions: The Shuttle (2008); and Ocean oasis (2000) (see also a recent study by Takács et al., 2019). The films were presented in a counterbalanced order across participants. A few emotionally arousing scenes were cut from the documentaries without creating strange transitions in the narrative actions. The final versions of the documentaries were mood-neutral and mentally non-demanding.

### Procedure in the fatigue group

The study meets ethical standards according to the Declaration of Helsinki and was approved by the Ethics Committee of the University of Pécs (nr. 7698). Figure 2 schematizes the procedure of the experiment in the two groups. Participants were asked to get adequate sleep during the night prior to the experiment and to abstain from alcohol and caffeine-containing substances before the experiment. In addition, they were told that they should avoid exhausting physical and mental activities (e.g. physical workout, studying for a class) before the experiment. Participants’ sleep duration was monitored using an actigraph (Philips Respironics Actiwatch-2; fatigue experiment: 7.46h, SD = 1.64h; control experiment: 7.82h, SD = 1.48h) and by self-reporting (fatigue experiment: 7.67h, SD = 1.61); control experiment: 7.82h, SD = 1.48). Participants in the two experiments did not significantly differ in the self-reported sleep (t(39) = 1.15, *p* = .26) or in the actigraph data (t(39) = − .74, *p* = .46).

**Figure 2.**
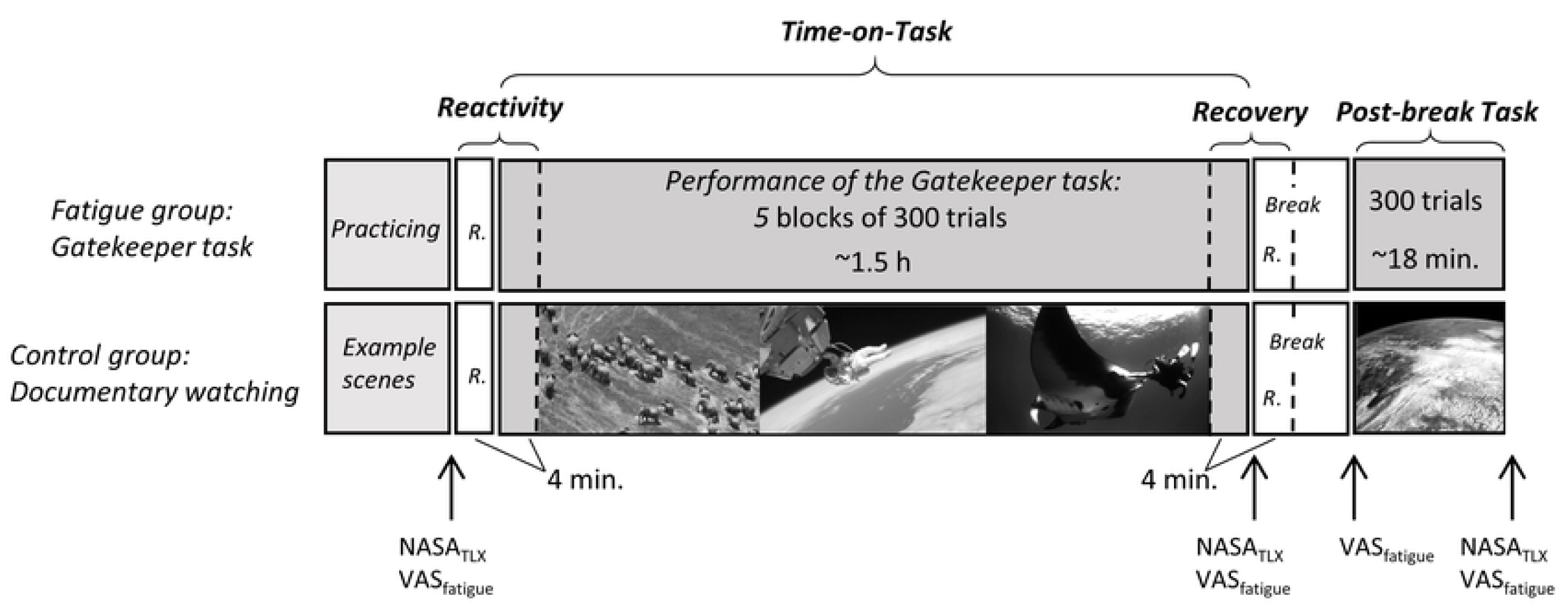
The schematized procedure of the study in the fatigue and the control (documentary watching) group. Participants in both groups had a 4-minute resting period before the time-on-task phase when ECG was recorded with eyes open and without body movements. It was followed by the time-on-task phase, when the participants in the fatigue group performed *5* blocks of 300 trials of the gatekeeper task without rest (each block lasted about 18 minutes). The participants in the control group watched documentaries without rest (5 blocks of 18 minutes). ECG was continuously recorded during the time-on-task phase in both groups. To calculate HRV within the experimental blocks, we selected the middle 15 minutes in each block. After that, participants had a break of 12 minutes. During the break, a 4-minute ECG was recorded again. Then, an additional block of 300 trials (appx. 18 minutes) were administered for the fatigue group and documentaries were presented for the control group. R: resting ECG. Participants indicated their subjective fatigue and workload multiple times during the experiment.

The experimental sessions started between 9:30 a.m. and 13:30 p.m. Participants were seated in a soundproof and uniformly air-conditioned (23C°) laboratory. Following the explanation of the cognitive task, three electrocardiographic (ECG) chest electrodes (Lead II.) were set up. In the fatigue group, the participants performed *72* practice trials. None of the participants indicated sensory irritability or unpleasantness after the practice trials. In addition, participants filled in two scales regarding the subjective task load and fatigue (see Subjective Measures section).

After the completion of the self-reported measures, a *4*-min-long resting ECG was recorded (with eyes open). Then, the prolonged performance of the task followed. During this ToT phase of the gatekeeper task the participants performed *5* blocks of 300 trials without rest. The exact duration of this phase depended on the participants’ response time, and thus slightly varied across participants (average duration total time-on-task: 1.48 h, range: 1.31 – 1.67 hours, *SD* = 0.1 h). When the 5 blocks of trials were completed, participants again filled in the subjective measures. Subsequently, participants had a break of 12 minutes. During the first 4 minutes of this break period, a resting ECG was recorded. After the break, participants were asked again to indicate their fatigue level. Then, an additional block of 300 trials were administered, after which the participants again had to fill in their fatigue level and perceived load during the last block.

### Procedure in the control group

In the control experiment, the procedure was identical to that of the fatigue experiment. Participants first had a familiarization period by being shown some example scenes from the documentaries (these scenes were not presented later in the prolonged viewing period). Participants were then instructed to watch the films, but it was emphasized that they did not have a particular task to perform (e.g. they did not need to memorize information provided by the documentaries). The ECG recording and the administration of the subjective measures (fatigue, task load) followed the same procedure as in the fatigue experiment. The only difference was that we reduced the number of scales used to assess participants’ subjective workload in the control experiment: participants reported only the perceived mental and physical demand and their frustration level. When the prolonged film-watching period ended, we asked the participants to rate how interesting (on a scale running from 1 to 7, where point 7 indicated an extremely interesting film), and emotional (on a scale running from −3 to +3, where 0 indicated an emotionally completely neutral film) the film was. Participants indicated that the film was moderately interesting (*M* = 5.14, range: 3-7, *SD* = 1.27) and only minimally emotionally arousing (*M* = 1.00, range: -1-3, *SD* = 1.05).

### Subjective measures of fatigue and workload

In each of the two experiments, participants completed the NASA Task Load Inventory (NASA_TLX_; Hart and Staveland, 1988) three times: after the practice (i.e. in the control group, after watching examples documentary scenes); after ToT period; and after the post-break task. The NASA_TLX_ is a multidimensional self-reported measure to assess individuals’ perceived workload during the task on *6* scales with 21 gradations: mental demand; physical demand; temporal demand; overall performance level; effort; and frustration level. Participants were also asked to indicate the level of fatigue they experienced on a Visual Analogue Scale (VAS_fatigue_ I.; 100mm long line, “No fatigue at all” was printed on the left side and “Very severe fatigue” on the right side).

#### Heart rate variability measurement

*ECG* data were digitized at a sampling rate of 1 kHz at 16-bit resolution with a CED 1401 Micro II analogue-digital converter device (CED, Cambridge, UK). The ECG signals were first visually inspected, and artefacts were corrected, and if necessary removed. Subsequently, participants’ R-R intervals, in milliseconds, were extracted using Spike2 software. The time elapsed between two successive R-waves (R-R intervals) were analyzed further by Kubios HRV analysis package 2.0 (Tarvainen et al., 2014). The artefacts within the R-R intervals were again corrected using the low artefact correction option of the Kubios software: detected artefact beats were replaced using cubic spline interpolation. Frequency-domain, time-domain, and non-linear HRV measures were calculated.

The frequency indices included the absolute high frequency power (0.15Hz - 0.4 Hz; ms^2^; HF), the log-transformed high frequency power (_ln_HF), the absolute low frequency power (0.04Hz – 0.15Hz; ms^2^; LF), and the log-transformed low frequency power (_ln_LF). We also used the powers of HF and LF in normalized units: the relative value of each power component in proportion to the total power minus the very low frequency (VLF, 0.003-0.04 Hz) component (i.e. LF_nu_ = (LF / total power – VLF) x 100; HF_nu_ = (HF / total power – VLF) x 100). In addition, the ratio between LF and HF band powers (LF / HF ratio) was calculated. We report the analyses of normalized units and the LF / HF ratio because they are frequently reported indices in HRV research, but it is important to note that recently the interpretability of these indices has been seriously challenged.

The time-domain measures included the mean heart rate (HR, beats/min), the root mean square of successive differences (RMSSD, ms), the log-transformed RMSSD (_ln_RMSSD), and the percent of the number of pairs of adjacent R-R intervals differing by more than 50 ms (pNN50; %).

The non-linear measures included sample entropy for quantifying signal complexity as well as the short-term HRV as a measure of the width of the Poincaré cloud (SD1), and the long term HRV as a measure of the length of the Poincaré cloud (SD2). Nevertheless, Ciccone et al., (2017) demonstrated that RMSSD and SD1 are both mathematically and empirically identical indices of HRV (i.e. SD1 equals to RMSSD multiplied by 1/ √2). In line with this, each analysis in the current study returned identical results for RMSSD and SD1 up to the third decimal. Therefore, below we do not report the results for SD1.

Recently, there has been a growing number of studies suggesting that HRV should be normalized with respect to average heart rate (Sacha, 2014; Sacha and Pluta, 2008; Sacha, 2013; Quintana and Heathers, 2014). Therefore, we also calculated and analyzed R-R normalized HRV indices. These analyses provided the same conclusions as those without RR normalization. The methods and the results of the R-R normalized analyses are shown in the supplementary materials (see Table S6).

We used two different intervals for the calculation of each HRV index: 4-minute intervals; and 15-minute intervals. The 4-min intervals were the resting period before the experiment, the first 4 minutes of the first experimental block, the last 4 minutes of the fifth experimental block, the resting period during the break, and, finally, the first 4 minutes of the post-break task block. These short intervals were used in the analysis of the reactivity and recovery effects (see Data Analysis section below). Studies addressing reactivity and recovery related changes in HRV often use even shorter intervals (e.g. Fiol-Veny et al., 2019; Kaczmarek et al., 2019; Feda and Roemmich, 2016). To calculate HRV within the experimental blocks, we selected the middle 15 minutes in each block. Please note that time-on-task blocks lasted about 18 minutes each but were not completely identical in terms of duration which depended on the participants’ reaction time. We also calculated and analyzed HR and HRV indices in the time-on-task with variable block intervals (i.e. full-length blocks), and these results are shown in the Supplementary materials (see Table S5). The conclusions are identical regardless of whether the analyses were performed for identical intervals (15-min) or for the full-length blocks.

In the resting period before the experiments, only sample entropy was significantly different between the two groups: it was significantly higher in the fatigue group than in the control group (*p* < .05). Heart rate (HR) and all the other HRV variables were not significantly different between the two groups (*p* = .09 - .94).

In addition to the HRV measures, for each trial, post-response cardiac activity was also calculated as the average difference in the R-R intervals during the 2.5s-long post-response period. The larger average difference reflected a slower activity after response.

#### Data Analysis

Statistical analysis was performed by JASP Version 0.11.1.0 (https://jasp-stats.org/). Changes in subjective fatigue (i.e. VAS_fatigue_) were analyzed by repeated measures analysis of variance (*r*ANOVA) with Administration Time (the 4 administrations of VAS_fatigue_) as a within-subject factor and Group as a between-subject factor. For the analysis of subjective workload (i.e. NASA_TLX_), the *r*ANOVA was performed with the within-subject factors of Administration Time (the 3 administrations of the NASA_TLX_), and Scales (the 3 scales of NASA_TLX_) and with Group as a between-subject factor.

For the performance measures of the gatekeeper task, the *r*ANOVA included Block (5 blocks of trials) and Target types as within-subject factors. Indicators of performance were Accuracy, Reaction time on correct responses (RT), Target-sensitivity (i.e. *d*’; Z_hit_ – Z_false alam_), and combined score of *d*’ and RT (Z_d’_ – Z_RT_). Separate ANOVAs were performed to analyze the ToT (changes from block 1 to block 5), and the break-related effects (changes from block 5 to block 6).

The reactivity of HRV was tested by *r*ANOVA (Block as within-subject factor, and Group as between-subject factor) comparing the 4-minute resting period data with the HRV in the first 4 minutes of the first experimental block. ToT-related HRV was analyzed including HRV data from the first to the fifth block (i.e. the 15-minute intervals selected in each of the blocks). Recovery-related HRV was tested by comparing the last 4 minutes of the fifth block and the first 4 minutes of the break period. Finally, the post-break reactivity in HRV was analyzed by a comparison of the HRV of the post-experiment resting period with the first 4 minutes of the post-break task block. In post-response cardiac activity analysis, we compared the correct and incorrect trials as a function of ToT. In each analysis Greenhouse-Geisser (ε) adjustment was applied if sphericity was violated. Bonferroni correction was applied to all follow-up analyses (non-planned) of the significant main effects and interactions.

## Results

### Subjective fatigue and workload

The two groups did not significantly differ in fatigue before the experiment (t(39) = −1.51, *p* = 0.14). The post-hoc analyses of the Administration Time x Group interaction (F(1,39) = 14.40, *p* < .01, η_*p*_^2^ = .27) indicated, however, that compared to the control group, participants in the fatigue group became more fatigued by the end of the ToT (fatigue group: *p* < .001; control group: *p* = 1.0). Within each group, subjective fatigue was not significantly affected by the break and showed no further change in the post-break performance block. For the workload measures, the Administration Time x Scales x Group interaction was significant (F(4, 36) = 5.91, *p* < .01, η_p_^2^ = .40). Post-hoc analyses indicated a significant ToT-related increment in the fatigue group (Mental demand: *p* < .001, Frustration: *p* = .01 Physical demand: *p* < .001), but no significant change in the control group (*p*s: .49 – 1.0). To summarize, the analyses of subjective fatigue and workload data suggest that the fatigue and workload manipulation was successful. Descriptive statistics for subjective fatigue and workload are presented in the supplementary materials (Table S1).

### Cognitive performance in the gatekeeper task

The main results of the cognitive performance in the gatekeeper task are reported in Table 1, and depicted in Figure 3 (descriptive statistics are in the supplementary materials, Table S2). The main effect of Block (1 to 5) reached significance for RT, and the composite scores. Post-hoc analyses revealed that participants’ RT became faster and their sensitivity to the targets increased from the first to the third block, but showed no significant change thereafter (for both RT and composite scores, block 1 vs. block 3: *p* < .01; block 3 vs block 5: *p =* 1.0). Accordingly, the findings suggest that, despite the strong increase in subjective fatigue and task load, there was no direct decline in performance on the gatekeeper task. Note, however, that we did find a highly significant improvement in performance after the break, suggesting that performance before the break may nevertheless have been suboptimal. The improvement was especially strong on *d*’ and the composite scores (see Figure 3).

**Table 1.** Results of *r*ANOVAs for four performance measures in the fatigue group.

| Variables | Analysis |  |  |  |  |  |
| --- | --- | --- | --- | --- | --- | --- |
|  | time-on-task (block 1-to-block 5) |  |  | break-related effects (block 5-to-block 6) |  |  |
|  | Block effect | Target effect | Block x Target | Block effect | Target effect | Block x Target |
| *RT* |  |  |  |  |  |  |
| df | 4,76 | 3,57 | 12,228 | 1,19 | 3,57 | 3,57 |
| F | 18.89*** | 66.14*** | 1.78 | 2.71 | 32.69*** | .06 |
| $\eta_p^2$ | .50 | .78 | .09 | .12 | .63 | .00 |
| *Accuracy* |  |  |  |  |  |  |
| df | 4,76 | 3,57 | 12,228 | 1,19 | 3,57 | 3,57 |
| F | 1.45 | 26.78*** | 2.11* | 13.09** | 10.25*** | 2.01 |
| $\eta_p^2$ | .07 | .58 | .10 | .41 | .35 | .10 |
| *d'* |  |  |  |  |  |  |
| df | 4,76 | 2,38 | 8,152 | 4,76 | 2,38 | 8,152 |
| F | 2.22 | 29.73*** | 2.26<sup>m</sup> | 14.72** | 22.25*** | 2.59 |
| $\eta_p^2$ | .10 | .61 | .11 | .44 | .54 | .12 |
| *Z<sub>d'</sub> - Z<sub>RT</sub>* |  |  |  |  |  |  |
| df | 4,76 | 2,38 | 8,152 | 4,76 | 2,38 | 8,152 |
| F | 9.57*** | 59.12*** | 1.96<sup>m</sup> | 10.74** | 37.87*** | 1.08 |
| $\eta_p^2$ | .33 | .76 | .10 | .36 | .66 | .05 |

Note: \* $p < .05$ , \*\* $p < .01$ , \*\*\* $p < .001$ , m: $p = .05$ ; $Z_{d'} - Z_{RT}$ : Combined score; $d'$ : target sensitivity

**Figure 3.**
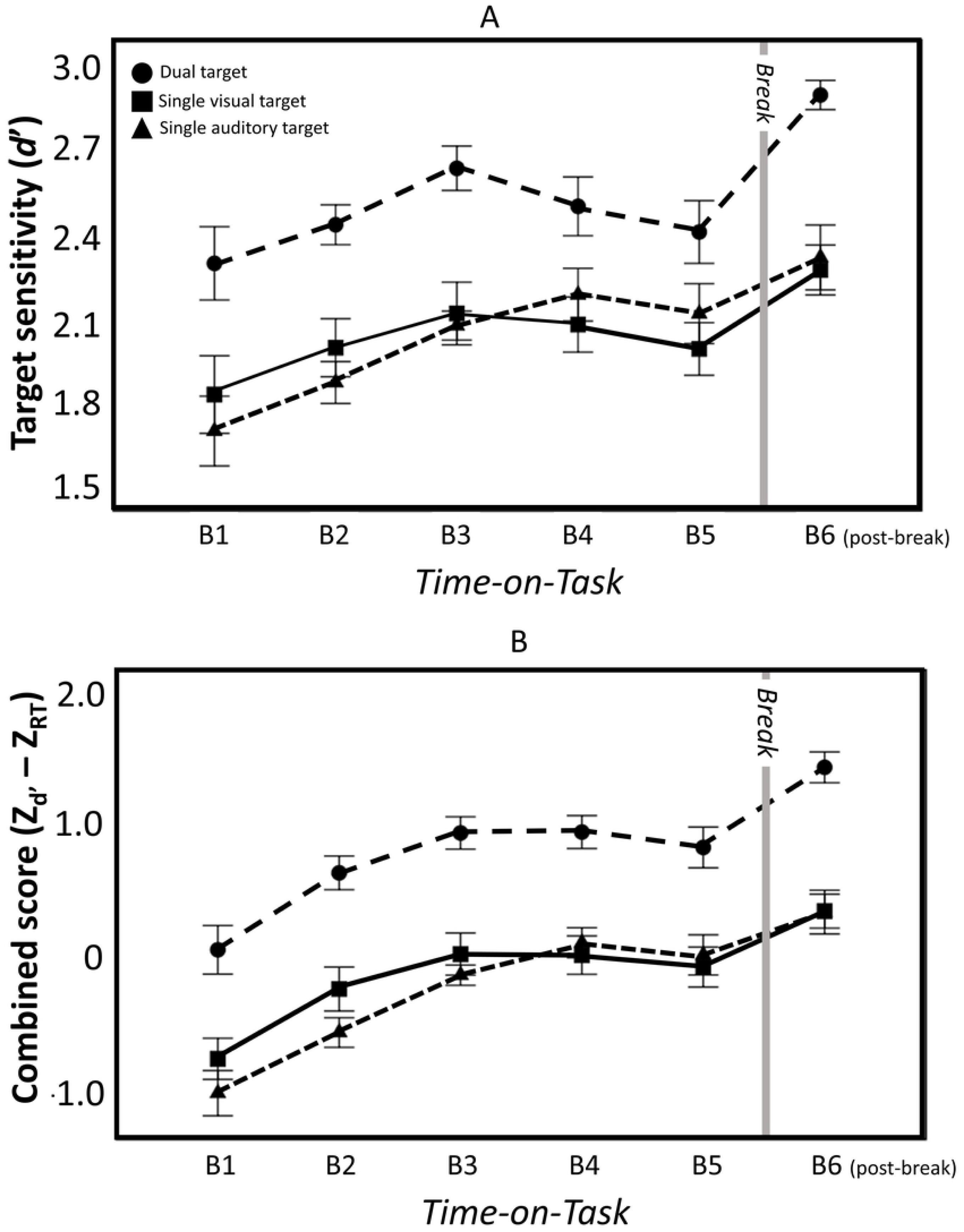
Results of 2 performance measures in the gatekeeper task (fatigue group): (A) Target sensitivity (*d*’), (B) Composite score (Z_d’_ – Z_RT_). Separate ANOVAs were performed to analyze the time-on-task (changes from block 1 to block 5), and the break-related effects (changes from block 5 to block 6). For dual target condition (circle), both the visual and auditory stimuli were identical to those presented two trials earlier. For the single auditory target condition (triangle), a 2-back match occurred for the auditory stimulus. For the single visual target condition (square), a 2-back match occurred for the visual stimulus. Error bars represent the within-subject error bars (Cousineau, 2005).

### Changes in HR and HRV in reactivity

Reactivity analyses focused on the changes in HR and HRV from the resting interval to the first 4 minutes of block 1. Table 2 presents the results of the analyses, and Figure 4 depicts the results for the HR and for HRV indices from each domain (for descriptive statistics see Table S3 and S4 in the supplementary materials). The *r*ANOVAs showed significant Interval x Group interactions for HR, _ln_LF, _ln_RMSSD, and _ln_HF. Post-hoc analysis revealed that, compared to the rest interval HR significantly increased during the first 4 minutes of the gatekeeper task (*p* < .001) in the fatigue group, but, in the control group, HR did not significantly change. In fact, in the control group there was a tendency (*p* = .08) towards a decreased HR. _ln_LF in the fatigue group significantly decreased in the first 4 minutes of the task (*p* < .001), but not in the control group (*p* = 1.00). Post-hoc analyses showed that there was no significant change from rest to the first 4 four minutes in _ln_RMSSD and _ln_HF, in either of the groups. This latter finding suggests that reactivity was not associated with a significant change in vagal activity.

**Table 2.** Results of *r*ANOVAs for the changes in Heart Rate and HRV in Reactivity.

| Variables | Analysis |  |  |  |
| --- | --- | --- | --- | --- |
|  | Block effect |  | Block x Group |  |
| | $F_{(1,39)}$ | $\eta_p^2$ | $F_{(1,39)}$ | $\eta_p^2$ |
| HR | 7.09* | .15 | 39.52*** | .50 |
| RMSSD | .01 | .00 | 1.98 | .05 |
| $\ln$RMSSD | 1.94 | .05 | 5.65* | .13 |
| pNN50 | .41 | .01 | 3.14<sup>m</sup> | .08 |
| HF | 2.24 | .05 | 1.04 | .03 |
| $\ln$HF | .00 | .00 | 6.25* | .14 |
| HFnu | 20.24*** | .34 | 1.02 | .03 |
| LF | 5.57* | .13 | .06 | .00 |
| $\ln$LF | 17.08*** | .31 | 11.46** | .23 |
| LFnu | 20.36*** | .34 | 1.04 | .03 |
| LF/HF | 14.60*** | .27 | .02 | .00 |
| SampEN | 9.66** | .20 | .89 | .02 |
| SD2 | 19.06*** | .33 | 3.16 | .08 |

Note. $r$ ANOVA: Block as within, and Group as between subject factors
Block: the 4-minute-long resting period before the experiment and the first 4 minutes of the first experimental block.
Group: group of participants in the fatigue and in the control experiment
\* $p < .05$ , \*\* $p < .01$ , \*\*\* $p < .001$ , m: $p = 0.09$

**Figure 4.**
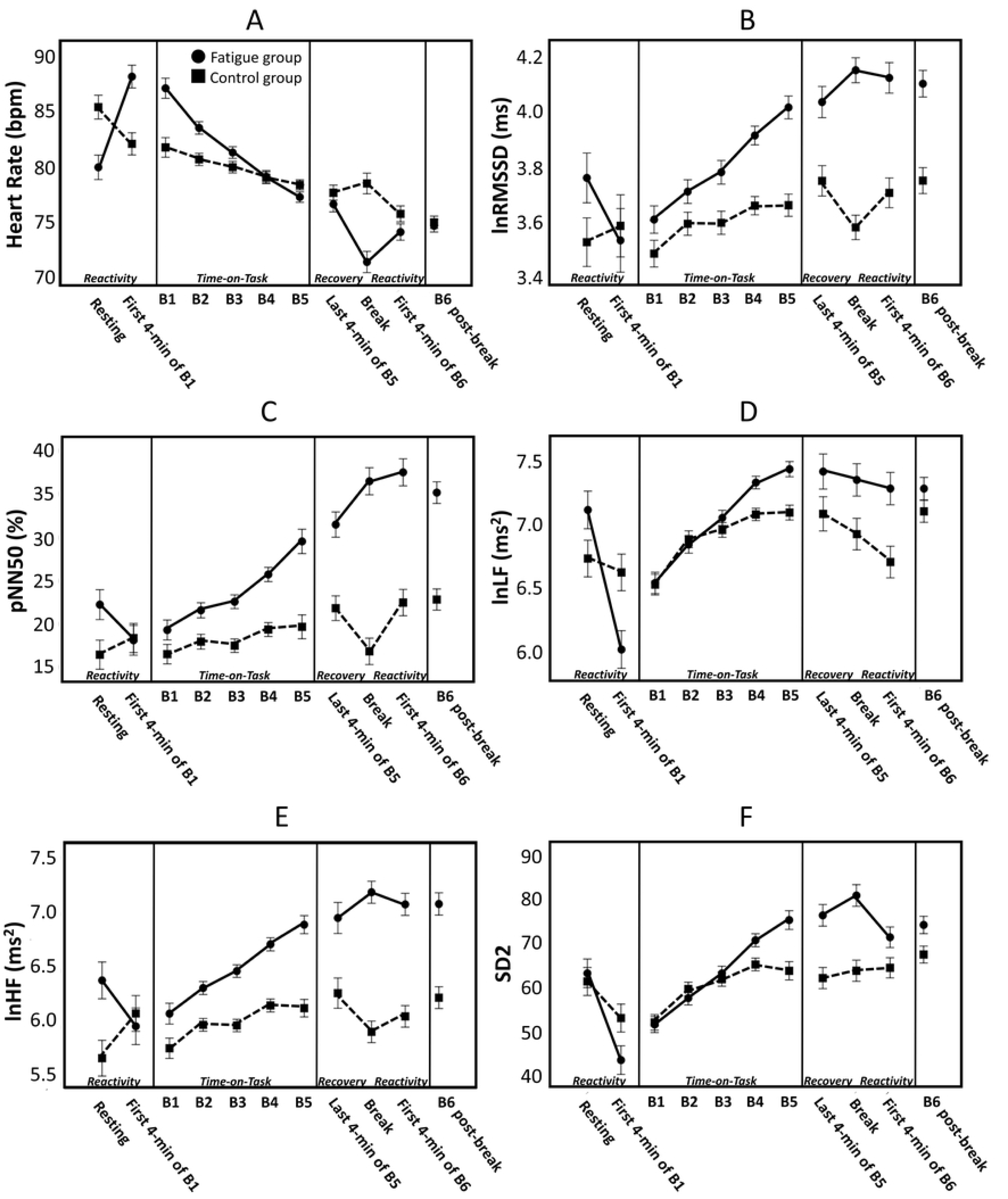
Results of mean heart rate (A) and five heart rate variability measures (B-F) in the fatigue (circle) and the control (square) group. Reactivity, time-on-task, and recovery related HRV was tested with separate ANOVAs. The reactivity of HRV was tested by *r*ANOVA comparing the 4-minute resting period data with the HRV in the first 4 minutes of the first experimental block. Time-on-task-related HRV was analyzed including HRV data from the first to the fifth block (i.e. the 15-minute intervals selected in each of the blocks). Recovery-related HRV was tested comparing the last 4 minutes of the fifth block and the first 4 minutes of the break period. The post-break reactivity in HRV was analyzed by a comparison of the HRV of the post-experiment resting period with the first 4 minutes of the post-break task block. The figure presents the results for HRV measures with the most robust Block x Group interaction. Error bars represent the within-subject error bars (Cousineau, 2005). _ln_RMSSD: the log-transformed root mean square of successive differences; pNN50: percent of the number of pairs of adjacent R-R intervals differing by more than 50 ms; lnLF; log-transformed Low frequency power; _ln_HF: the log-transformed High frequency power; SD2: the length of the Poincaré cloud.

### Changes in HR and HRV with time-on-task

The time-on-task analyses focused on the changes in HR and HRV from block 1 to block 5 (see Table 3). The significant Block x Group interactions showed that the following cardiac parameters changed differentially in the fatigue and the control groups: HR; RMSSD; _ln_RMSSD; pNN50; HF; _ln_HF; LF; _ln_LF; and SD2. Post-hoc analyses revealed that, compared to the control, there was a larger HR decrement in the fatigue group (block 5 vs block 1 in the fatigue group: *p* < .001, in the control group: *p* = 0.01). In the fatigue group, RMSSD, pNNF50, and HF (three vagal mediated HRV components) significantly increased from block 1 to block 5 (block 5 vs block 1 for RMSSD: *p* < .001, pNN50: *p* < .001, HF: *p* = .001), but no significant changes were observed in the control group. _ln_RMSSD, _ln_HF, LF, _ln_LF, and SD2 showed significant increases in the fatigue and the control group, but those changes were always larger in the fatigue group (see also Figure 4).

**Table 3.** Results of *r*ANOVAs for the changes in Heart Rate and HRV in time-on-task.

| Variables | Analysis |  |  |  |
| --- | --- | --- | --- | --- |
|  | Block effect |  | Block x Group |  |
| | $F_{(4,156)}$ | $\eta_p^2$ | $F_{(4,156)}$ | $\eta_p^2$ |
| HR | 42.06*** | .52 | 9.90*** | .20 |
| RMSSD | 10.91*** | .22 | 3.64* | .09 |
| lnRMSSD | 23.11*** | .37 | 4.39** | .10 |
| pNN50 | 13.03*** | .25 | 3.67* | .09 |
| HF | 3.58* | .08 | 2.84* | .07 |
| lnHF | 21.53*** | .36 | 3.23* | .08 |
| HF<sub>nu</sub> | 3.31* | .08 | .48 | .01 |
| LF | 17.71*** | .31 | 2.74<sup>m</sup> | .07 |
| lnLF | 41.61*** | .52 | 3.13* | .07 |
| LF<sub>nu</sub> | 3.31* | .08 | .48 | .01 |
| LF/HF | 1.58 | .04 | .35 | .01 |
| SampEN | 1.08 | .03 | 1.70 | .04 |
| SD2 | 35.11*** | .47 | 5.04** | .11 |

*Note.* *r*ANOVA: Block as within, and Group as between subject factors
Block: five 15-minute-long intervals (selected in each block) in the Time-on-Task phase.
Group: group of participants in the fatigue and in the control experiment
\* $p < .05$ , \*\* $p < .01$ , \*\*\* $p < .001$ , m: $p = 0.05$

In addition, we analyzed the association between ToT-related changes in subjective fatigue (i.e. the difference between fatigue after ToT and before) and HRV (i.e. the difference between block 5 and block 1): a higher increment in HRV was found to be significantly or marginally significantly associated with an enhanced level of subjective fatigue (RMSSD: r_s_ = .42, *p* = .07; pNN50: r_s_ = .39, *p* = .09; HF: r_s_ = .45, *p* = .04; LF: r_s_ = .74, *p* < .001; SD2: r_s_ = .57, *p* = .009).

### Changes in HR and HRV during recovery

Recovery analyses involved the changes in HR and HRV from the last 4 minutes of block 5 to the 4-minute-long resting period in the break (see Table 4). We found that, during the recovery period the following measures changed differentially in the two groups: HR; RMSSD; _ln_RMSSD; pNN50; _ln_HF; HF_nu_; and LF_nu_. While participants’ HR in the fatigue group significantly decreased (*p* < .001) during the recovery period, no significant change was found in the control group. In contrast for _ln_HF, HF_nu_, and LF_nu_, the post-hoc analyses did not reveal a significant difference between the last 4 minutes of block 5 and the break.

**Table 4.** Results of *r*ANOVAs for the changes in Heart Rate and HRV in Recovery.

| Variables | Analysis |  |  |  |
| --- | --- | --- | --- | --- |
|  | Block effect |  | Block x Group |  |
| | $F_{(1,39)}$ | $\eta_p^2$ | $F_{(1,39)}$ | $\eta_p^2$ |
| HR | 8.85** | .19 | 16.74*** | .30 |
| RMSSD | .24 | .01 | 4.75* | .11 |
| lnRMSSD | .27 | .01 | 7.69** | .17 |
| pNN50 | .00 | .00 | 12.00** | .24 |
| HF | 1.71 | .04 | 1.00 | .03 |
| lnHF | .18 | .01 | 4.42* | .10 |
| HF<sub>nu</sub> | .13 | .00 | 7.48** | .16 |
| LF | .08 | .00 | 2.74 | .07 |
| lnLF | .68 | .02 | .12 | .00 |
| LF<sub>nu</sub> | .13 | .00 | 7.52** | .16 |
| LF/HF | .01 | .00 | .91 | .02 |
| SampEN | .78 | .02 | .04 | .00 |
| SD2 | 1.62 | .04 | .34 | .01 |

*Note.* *r*ANOVA: Block as within, and Group as between subject factors
Block: the last 4 minutes of block 5 and the 4-minute-long resting period in the break
Group: group of participants in the fatigue and in the control experiment
\* $p < .05$ , \*\* $p < .01$ , \*\*\* $p < .001$

### Reactivity in HR and HRV after the break

Reactivity after the break was analyzed by comparing the first 4 minutes of block 6 with the 4-minute resting period during the break (see Table 5). There was a significant Block x Group interaction for 4 cardiac parameters: HR; RMSSD; HF_nu_; and LF_nu_. Post-hoc analyses revealed that while participants’ HR significantly increased in the fatigue group after the break (*p* < .01), it had a significantly opposite, decreasing pattern in the control experiment (*p* < .01). HRV in the fatigue group remained about the same level as during the break. The control group tended to show a reactivity-related change, but this effect did not reach significance in the post-hoc analyses. These findings generally suggest that, similarly to reactivity at the beginning of experiment, reactivity after the break was not accompanied with increased vagal withdrawal (i.e. reduction in vagal control). In addition, the findings also clearly indicated that the marked improvement in the performance of the gatekeeper task after the break was not associated with concurrent changes in HR and HRV.

**Table 5.** Results of *r*ANOVAs for the changes in Heart Rate and HRV in Reactivity after the break.

| Variables | Analysis |  |  |  |
| --- | --- | --- | --- | --- |
|  | Block effect |  | Block x Group |  |
| | $F_{(1,39)}$ | $\eta_p^2$ | $F_{(1,39)}$ | $\eta_p^2$ |
| HR | .00 | .00 | 26.91*** | .41 |
| RMSSD | 2.01 | .05 | 4.68* | .11 |
| lnRMSSD | 1.53 | .04 | 3.75<sup>m</sup> | .09 |
| pNN50 | 4.66* | .11 | 2.22 | .05 |
| HF | .14 | .00 | 1.81 | .04 |
| lnHF | .02 | .00 | 1.60 | .04 |
| HFnu | 2.45 | .06 | 4.16* | .10 |
| LF | 1.16 | .03 | .29 | .01 |
| lnLF | 1.31 | .03 | .36 | .01 |
| LFnu | 2.52 | .06 | 4.20* | .10 |
| LF/HF | 3.03<sup>m</sup> | .07 | 2.65 | .06 |
| SampEN | 6.47* | .14 | 2.77 | .07 |
| SD2 | 2.52 | .06 | 3.33<sup>m</sup> | .08 |

*Note.* *r*ANOVA: Block as within, and Group as between subject factors
Block: the 4-minute-long resting period of the break and the first 4 minutes of block 6
Group: group of participants in the fatigue and in the control experiment
\* $p < .05$ , \*\* $p < .01$ , \*\*\* $p < .001$ , m: $p < 0.09$

### Motivation and inattention: Analysis of post-error cardiac activity, post-error reaction times, and reaction time variability

In the gatekeeper task, the analysis of phasic heart rate activity after responses yielded a significantly slower cardiac activity after inaccurate responses were compared to correct responses (F(1,19) = 9.65, *p* < .01, η_*p*_^2^ = .34). The Block x Correctness of Responses interaction, however, was non-significant suggesting that this error-related cardiac activity remained unchanged over the ToT (F(4,76) = 1.13, *p* = .35). Similarly, post-error slowing was significant: responses were generally slower in a trial if the response in the previous trial was erroneous (F(1,19) = 44.42, *p* < .001, η_*p*_^2^ = .70), but no significant interaction with ToT was obtained (F(4,76) = .3, *p* = .78). All in all, these findings seem to indicate that the participants did not strongly decrease in their willingness to do well on the task because their reaction to errors remains relatively stable over time.

Finally, RT variability (i.e. mean / SD) – as a frequently used index of the lapses in attention, or inattention level – was calculated for each block of trial. We found that RT variability increased during the prolonged performance period but this change did not reach significance, suggesting that participants did not become substantially inattentive during the prolonged performance of the task (F(4,76) = 1.13, *p* = .10., η_*p*_^2^ = .11).

## Discussion

The present study examined the temporal profile of HRV, including the changes related reactivity, ToT, and recovery. Our hypotheses were based on previous observations that prolonged performance on a cognitively demanding task is associated with reduced norepinephrine levels that coincide with elevated levels of subjective fatigue, decreased task-engagement, and compromised cognitive performance (Hopstaken et al., 2015a, 2015b, 2016). In line with this model, we expected to find ToT-related changes particularly on those HRV components that are presumed indices of vagal activity, and as such indirectly and partially reflect the activation of the LC-NE system (Laborde et al., 2017). We also analyzed post-response cardiac activity (i.e. heart rate change) as an autonomic measure of performance monitoring.

### Reactivity

In the reactivity period, the vagal-mediated HRV measures showed no significant decrement. This is line with our previous within-subject study in which the vagal-mediated HRV components in a cognitively demanding task did not show reactivity effects (Matuz et al., 2019). A possible explanation for the weak vagal withdrawal is that because the gatekeeper task was very demanding, parasympathetic inhibition may have tempered vagal withdrawal to prevent too-high of a level of arousal, which would have probably interfered with an optimal task concentration. We found a significant decrement of _ln_LF, though, suggesting that this component of HRV is accompanied with enhanced mental effort when individuals start working on a demanding cognitive task. Note that the LF component is affected by sympathetic as well as parasympathetic outflows (e.g. Akselrod et al., 1981; Ng et al., 2009) and does not have a direct correlation with norepinephrine levels (Alvarenga et al., 2006; Baumert et al., 2009; Moak et al., 2007). Oscillation in LF power is suggested to mainly provide information about blood pressure control, such as baroreflex sensitivity (Goldstein et al., 2011; Rahman et al., 2011). Yasumasu et al. (2006) found that baroreceptor function was inhibited during cognitive task performance possibly to allow increases in cardiovascular activity when exerting mental effort. A similar inhibitory process might also the reason for the decreased _ln_LF in the reactivity period that we found in the current study. The _ln_LF reactivity effect, however, is not to be generalizable, because it did not occur in the second reactivity period after the break. The significant increase in HR that was found in the reactivity period was in line with the notion that voluntary modulated high-level mental effort involves widespread cortical and subcortical activation, which is associated with increasing heart rate (Khachouf et al., 2017).

### Time-on-task

Five of the HRV measures (i.e. RMSSD, _ln_RMSSD, pNN50, HF, _ln_HF) that are presumed indices of vagal activity showed a marked increase over time in the fatigue group (but not in the control group). This suggests an increased vagal inhibition on heart activity with increasing ToT. This finding supports our hypothesis and fits with previous research suggesting a declined activation of LC-NE system due to fatigue (Hopstaken, et al., 2015a; 2015b). ToT-related increases in HF and RMSSD were also associated with increases in subjective mental fatigue. Subjective fatigue as well as lower LC-NE system activation are both known to be accompanied with task disengagement (Aston-Jones and Cohen, 2005; Hopstaken et al., 2016). Therefore, the pattern of subjective and HRV findings seem to indicate that, over time, participants tended to disengage from the gatekeeper task. The literature shows that task disengagement, however, is a complex phenomenon including at least three domains: i) concentration, ii) task motivation, and iii) energetic arousal (Matthews et al., 2010). The concentration aspect of task disengagement may not have changed that much with ToT, because participants’ reaction time variability as an index of concentration level showed no significant ToT-related change (see e.g. Albaugh, et al., 2017). Similarly, there are also several reasons to believe that the deliberate motivation aspect of engagement was not strongly diminished. First, actual performance remained relatively high over the course of the five blocks suggesting that the participants still tried to do their best despite rising feelings of fatigue and workload. Second, participants’ heart rate decelerated, and their response slowed down after making an erroneous response. This suggests that participants were still aware of their inaccurate responses and were motivated to exert compensatory effort during the whole duration of the ToT period. Subsequently, the increased vagal-mediated HRV may be particularly indicative of a decrease in energetic arousal, the third aspect of task disengagement. Energetic arousal was proposed first by Thayer (1978), and it has been suggested that a high level of energetic arousal reflects the mobilization of physiological and cognitive capabilities (Matthews and Davies, 2001). Yoshino and Matsuoka (2011) found that higher level of energetic arousal affected the shifting of activation of the autonomic nervous system toward sympathetic dominance expressed by a lower level of vagal-mediated HRV. In accordance with this, the present findings suggest that it primarily the participants’ energetic arousal that declined during ToT.

With increasing time spent on the gatekeeper task, the low frequency indices and the non-linear SD2 component of HRV also increased. This may indicate a ToT-related increase in baroreceptor sensitivity and suggests a decline in cognitive effort (Reyes del Paso et al., 2004). It should be noted, however, that there are also evidences showing that LF is not a robust index of baroreceptor sensitivity (Martelli et al., 2014).

### Recovery and reactivity after the break

After the strong increment in HRV over ToT, we found no further increase in recovery, nor did we find any significant reactivity effects after the break. The relatively high vagal activation that was maintained in the break possibly is an adaptive physiological reaction in order to replenish self-regulatory resources and to cope with potential further stressors (e.g. Kimhy et al., 2013). In line with this, participants showed a rather strong cognitive improvement after the break which, however, was not accompanied by changes in HRV. Specifically, participants’ cardiac activity (both in HR and HRV) in the post-break block was approximately the same level as during the last block (block 5) of the ToT period. Yet in this last block of trials, performance was worse than during the block after the break. This suggests that the break-related improvement did not necessarily require extra metabolic support in the form of increased activation of the cardiac system.

### Non-predictive measures and the importance of the control condition

Many HRV measures we calculated were found to be non-, or hardly sensitive to the different phases of the study including the normalized units of LF and HF as well as the LF / HF ratio. The interpretation of the lack of significant findings for these measures is compromised, however, due to their unclear physiological basis. For example, many studies now suggest that LF/HF should no longer be interpreted as an index of sympatho-vagal balance (e.g. Billman, 2013). Similarly, the interpretation of LF_nu_ and HF_nu_ is unclear because their physiological basis is questioned (Pagani, Lucini and Porta, 2012). As Heathers (2014) has argued, they are not reliable measurements if the very low frequency component is recorded in the short term only (i.e. fewer than 15 minutes; see, for example, the reactivity and recovery periods in the current study).

Finally and importantly, many of the HRV components we measured showed clear changes both in the fatigue and the non-fatigue groups (see e.g. _ln_RMSSD, _ln_HF, LF, _ln_LF, SD2) suggesting that HRV can also be significantly changed even if individuals are engaged in a prolonged non-fatiguing activity.

### Limitations of the study

Although the present study provides details of the changes in HR and HRV when engaging in a cognitively demanding task, the study also has at least two limitations to consider. First, in reactivity and recovery, a longer monitoring interval for the calculation of the time domain HRV indices would have been better. On the other hand, we aimed to specifically observe the period when individuals experience a change in cognitive demand, and therefore a longer interval would have extended beyond this psychologically special period. As mentioned above, HRV studies frequently use short monitoring intervals to observe changes related to reactivity and recovery (see e.g. Fiol-Veny et al., 2019; Kaczmarek et al., 2019). Second, the explanations about the underlying physiological source of HRV findings would have been benefitted from the monitoring of additional physiological measurements with clear physiological sources (e.g. pupillography, and skin conductance).

### Summary and Conclusion

To summarize the main findings of the present study, we showed that ToT on a dual 2-back task with a game-like character (i.e. the gatekeeper task) was associated with an elevated level of subjective fatigue and, concurrently, with decreased heart rate as well as increased HRV. Compared to a non-fatiguing documentary viewing experiment, the vagal-mediated HRV components showed a clear differential trend. Because higher LC activity has been found to be associated with lower parasympathetic influence on HRV (e.g. Mather et al, 2017), the present findings indirectly support the notion of the LC-NE involvement in mental fatigue. Increased levels of fatigue may be associated with a lowered activity of the LC-NE system. Based on post-error cardiac slowing and post-error RT analyses, we found no evidence for motivation deficit in association with increasing ToT. In addition, we found that – in contrast to HRV – HR tended to be changed in all three phases of the study: HR increased in reactivity, and decreased in time-on-task, and recovery.

## Author Contributions

András Matuz, *roles*: Conceptualization, Data Curation, Formal Analysis, Investigation Methodology, Writing – Original Draft Preparation

Dimitri van der Linden, *roles*: Conceptualization, Writing – Original Draft Preparation

Zsolt Kisander, *roles*: Formal Analysis, Methodology

István Hernádi, *roles*: Writing – Review & Editing, Funding Acquisition

Karádi Kázmér, *roles*: Writing – Review & Editing

Árpád Csathó, *roles*: Conceptualization, Data Curation, Formal Analysis, Funding Acquisition, Investigation, Methodology, Writing – Original Draft Preparation

## Data Availability

Data repository: https://data.mendeley.com/datasets/b3svkcpm5d/draft?a=4ba67d69-f985-4a93-b4ac-a9a602b8ff59

